# The *Nopp140* gene in *Drosophila melanogaster* displays length polymorphisms in its large repetitive second exon

**DOI:** 10.1101/513978

**Authors:** Sonu Shrestha Baral, Patrick J. DiMario

## Abstract

Nopp140, often called the nucleolar and Cajal body phosphoprotein (NOLC1), is a predicted chaperone for the transcription and processing of rRNA as it assembles into small and large ribosomal subunits. Nopp140 is conserved among the metazoans with an amino terminal LisH dimerization domain followed by a large central domain consisting of alternating acidic and basic motifs of low sequence complexity, and then a carboxyl terminus that can be variable due to alternative splicing. Treacle is another nucleolar chaperon in vertebrates that is structurally and functionally related to Nopp140. While the large central domains of vertebrate Nopp140 and treacle are encoded by several exons, the similar domain in *Drosophila* Nopp140 is encoded by a single exon. We define three overlapping repeat sequence patterns (P, P’, and P’’) within the central domain of *D. melanogaster* Nopp140. These repeat patterns are poorly conserved in other *Drosophila* species. A length polymorphism for the P’ pattern in *D. melanogaster* displays either two or three 96 base pair repeats, respectively referred to as *Nopp140-Short* and *Nopp140-Long*. PCR characterization of the long and short alleles shows a poorly-defined, artefactual bias toward amplifying the long allele over the short allele. Fly lines homozygous for one or the other allele, or heterozygous for both alleles, show no discernible phenotypes. The significance of this polymorphism lies in discerning the properties of the Nopp140’s large central domain in assemblage and phase separations in nucleoli and Cajal bodies.

## Introduction

The nucleolar phosphoprotein of 140 kDa (Nopp140) is a ribosome assembly factor that locates within the dense fibrillar component of nucleoli and within nuclear Cajal bodies (CBs) (Meier and Blobel, 1990, 1992). It was first described as a chaperone that shuttles between the nucleolus and cytoplasm (Meier and Blobel, 1992), perhaps to facilitate the import of other ribosome assembly factors that have SV40 T antigen-type nuclear localization signals (Meier and Blobel, 1990). Subsequent work showed that Nopp140 associates with C/D box small nucleolar ribonucleoprotein (snoRNPs) that guide nucleotide-specific 2′-*O*-methylation of the pre-rRNA, and with H/ACA box snoRNPs that direct site-specific pseudouridylation of pre-rRNA and snRNAs (Yang *et al.*, 2000). These interactions with Nopp140 may initiate within CBs as Nopp140 is thought to shuttle these snoRNPs from CBs to nucleoli (Yang *et al.*, 2000; Wang *et al.*, 2002; Meier, 2005; Lo *et al.*, 2006; He and DiMario, 2011). Nopp140 also interacts with RNA Pol I to regulate rRNA transcription, perhaps linking RNA Pol I transcription with pre-rRNA processing (Chen *et al.*, 1999; Meier, 2005).

The human Nopp140 homologue, called nucleolar and Cajal body phosphoprotein (NOLC1; CCDS 65925.1), is encoded by *NOLC1* (ID: 9221) which is located on chromosome 10 (NC_000010.11). *NOLC1* contains 13 exons that encode isoforms of 700 or 707 amino acids. Exon 1 encodes a Lis1 homology (LisH) domain that is likely used for dimerization (Mateja *et al.*, 2006). Exons 3-10 encode a large central peptide domain that consists of 10 repeating motifs; each motif contains a basic region rich in lysine, alanine, and proline followed by an acidic region rich in aspartate, glutamate, and phosphoserine. The serine residues are extensively phosphorylated *in vivo* by casein kinase type II (CKII) enzymes thus contributing to the acidic properties of the region (Meier, 1996; Li *et al.*, 1997). Six of the ten basic-acidic motifs in human NOCL1 are encoding by adjacent exons. The remaining four motifs are encoded by exon 10. Exons 11-13 in *NOCL1* encode the carboxyl tail of 119 amino acids with a conserved but putative protein kinase A phosphorylation site, indicating that Nopp140 is likely involved in regulatory signaling events (Meier, 1996). Because no mutations are known to exist for human *NOCL1*, *de novo* mutations in *NOCL1* are likely embryonic lethal.

Treacle is a nucleolar ribosome biogenesis factor related to Nopp140 in structure and function, but thus far found only in vertebrates. Human treacle is encoded by the *TCOF1* gene (gene ID: 6949; chromosome 5, NC_000005.10); haplo-insufficiency mutations in *TCOF1* result in the Treacher Collins syndrome (TCS), a ribosomopathy in which select neural crest cells undergo p53-dependent apoptosis resulting in impaired craniofacial development (Jones et al., 2008; Sakai and Trainor, 2009; Narla and Ebert, 2010). Similar to *NOLC1*, the *TCOF1* gene consists of multiple exons. Exon 1 again encodes a LisH domain, and similar to vertebrate Nopp140, treacle contains several repeating basic-acidic motifs comprising a large central domain (Wise et al. 1997). As with human *NOCL1*, multiple exons in the *TCOF1* gene encode these repeat peptide motifs (Isaac et al. 2000).

*Nopp140* is the closest gene in *Drosophila* spp. to *TCOF1* in humans. In *Drosophila melanogaster*, *Nopp140* maps to the left arm of chromosome 3 proximal to the centromere in cytological region 78F4. Two protein isoforms arise by alternative splicing. Nopp140-True contains 686 amino acids, while Nopp140-RGG contains 720 residues (Waggener and DiMario, 2002). Both isoforms localize to nucleoli and Cajal bodies when expressed exogenously as GFP fusions in transgenic embryos, larvae, and adults (McCain *et al.*, 2006). The mRNA encoding Nopp140-True consists of four exons: exon 1 encodes the LisH domain as in human NOCL1 and treacle. While the large central domains in human NOCL1 and treacle are encoded by several exons, the large central domain in *Drosophila* Nopp140-True consists of 16 repeating basic-acidic motifs, all encoded by exon 2. The carboxyl terminus of Nopp140-True is encoded by exons 3 and 4; this domain is 65% identical to the carboxyl terminus of human Nopp140 over a 94-amino acid stretch. Therefore, we consider Nopp140-True to be the *true* orthologue of vertebrate Nopp140 in *Drosophila* (Waggener and DiMario, 2002). The second isoform in *Drosophila*, Nopp140-RGG, is encoded by the same two exons 1 and 2, and is thus identical to Nopp140-True in its first 583 amino acid residues. Alternative splicing, however, generates a distinctly different carboxyl domain rich in glycine and arginine residues that form repeating RGG tri-peptide motifs (see Figure 1B). RGG motifs are common to many RNA-associated proteins (Raman et al., 2001).

**Figure 1.**
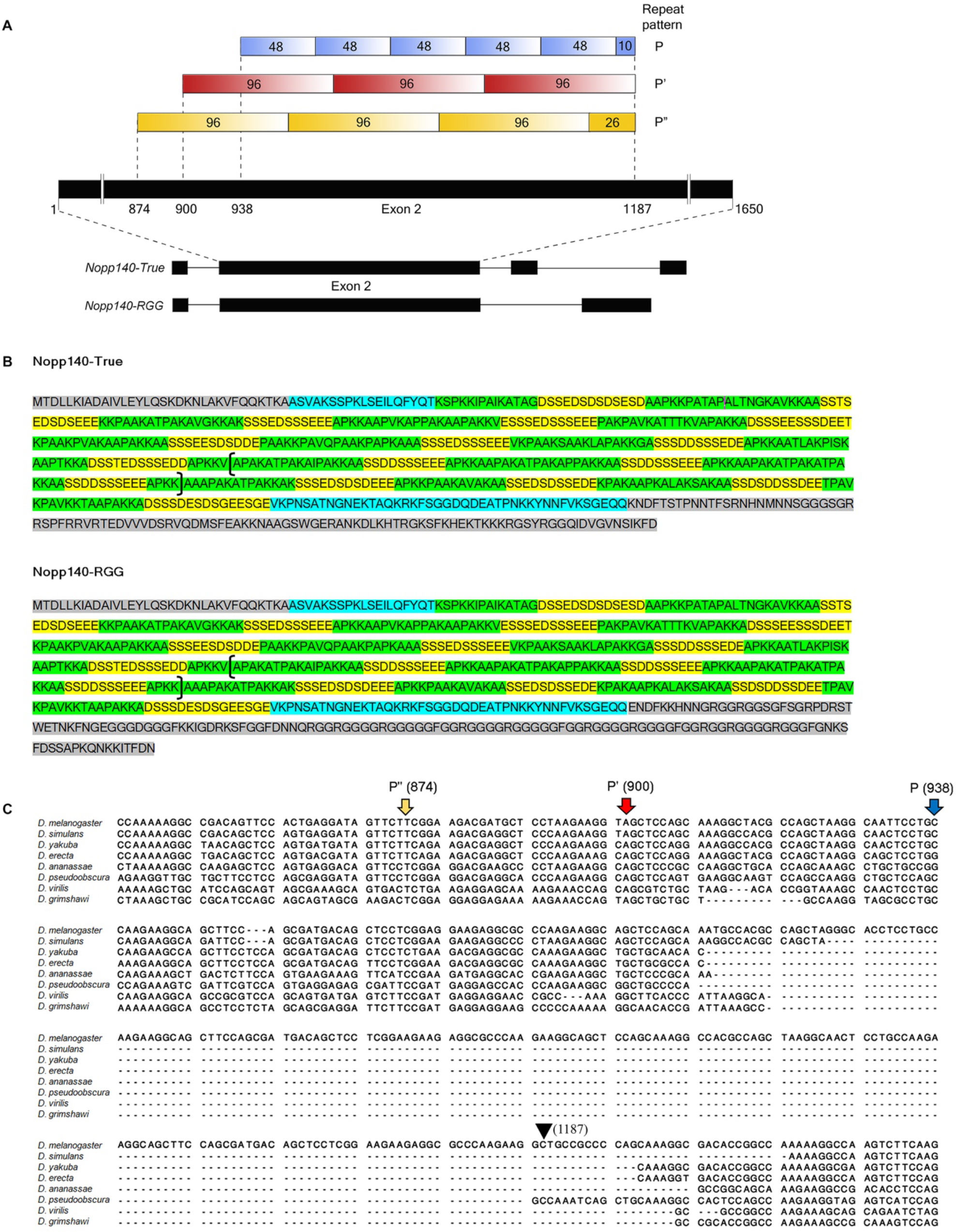
The second exon of *Nopp140* gene contains multiple overlapping repeat patterns unique to the *Drosophila melanogaster* species. **A** Schematic representation of a region within the second exon (1650 bp in length) of *Nopp140* gene that contains three repeat patterns: repeat pattern P with five 48 base bp repeats, repeat pattern P’ with three 96 bp repeats, and region P’’ with three 96 bp repeats different from that of pattern P’. *Nopp140* genomic regions for two isoforms of Nopp140 protein, Nopp140-True and Nopp140-RGG, are shown (solid blocks are exons and lines are introns). **B** Complete protein sequence of *Drosophila* Nopp140-True and Nopp140-RGG are shown. The beginning and ending sequence of the common second exon are highlighted in blue. Within the second exon, alternating acidic (containing aspartate and glutamate) and basic (containing lysine and arginine) motifs are highlighted in yellow and green respectively. The N-terminus (first exon which is common to both protein isoforms) and C-terminus (which is unique to each protein isoform) are highlighted in grey. The entire repeat pattern P’ region comprising of three 96 bp repeats is marked by brackets. **C** Sequence alignment of *Drosophila melanogaster Nopp140* gene and Nopp140-like protein coding gene from seven other *Drosophila* species. Shown here is the alignment of *Nopp140* second exon containing the three repeat patterns P, P’, and P’’. The first nucleotide of each repeat region is indicated by an arrow, the nucleotide number is in parenthesis (numbering begins at the first nucleotide of the second exon), and the arrowhead indicates the end of all repeats.

The large central domain of metazoan Nopp140 displays low amino acid complexity, and is intrinsically disordered, another common feature for RNA chaperones (Dyson and Wright, 2004; Tompa and Csermely, 2004). Here we describe several repeating but overlapping amino acid sequence patterns within the large central domain of Nopp140 in *Drosophila melanogaster*. We compare differences in these patterns within other *Drosophila* species, and we describe two *Nopp140* alleles in *Drosophila melanogaster*, *Nopp140-Long* and *Nopp140-Short*. The two alleles differ by exactly 96 bps that encode a 31 amino acid repeat unit within the central domain. Both alleles seem to be functional as there are no apparent adverse phenotypes in flies expressing one or the other allele. Polymorphisms in Nopp140’s central domain could potentially influence assemblage and phase separations (Toretsky and Wright, 2014) within nucleoli and CBs (Feric et al., 2016). Hence, the significance of this polymorphism lies in discerning the properties and function of the large central domain of metazoan Nopp140 orthologues.

## Materials and Methods

### Fly lines

Flies were obtained from the Bloomington *Drosophila* Stock Center (Indiana University) and maintained in the laboratory at room temperature on standard cornmeal medium. The following fly stocks were used: *w*^*1118*^(Bloomington stock 3605), *Daughterless-GAL4* (*Da-GAL4;* Bloomington stock 8641), *Oregon-R-C* (Bloomington stock 5), *Canton-S* (Bloomington stock 64349), *TM3/Et*^*50*^ (Bloomington stock 64349), *CyO/Sp*^*1*^ (Bloomington stock 4199), and *Nopp140* gene deletion line (He et al. 2015).

### Genomic DNA extraction

Healthy well-fed flies were frozen at −80°C for 10 minutes and subsequently homogenized in 100 mM Tris-HCl (pH 7.5), 100 mM EDTA, 100 mM NaCl, and 0.5% SDS. Following 30 min incubation at 70°C, genomic DNA was precipitated in 1:2 ratio of 5 M KOAc: 6 M LiCl on ice for 10 min. The precipitated genomic DNA was purified using phenol-chloroform extraction, followed by ethanol precipitation. The DNA pellet was suspended in deionized water, and the DNA concentration was measured by a NanoDrop 1000 Spectrophotometer.

### PCR amplification and Band analysis

PCR reactions were performed in 25 μL total volume containing 50-100 ng (unless otherwise specified) of genomic DNA, 0.40 μM of each primer (Table I), 0.160 mM of each dNTP, 0.30 mM of MgCl_2_, 0.5 X Phusion GC Buffer, and 0.40 unit of Phusion high-fidelity DNA polymerase (M0530S, New England BioLabs). Amplification was performed in a BIO-RAD C1000 Thermal Cycler with the following thermal cycling conditions: 2 min initial denaturation at 95°C, followed by 34 cycles of denaturation for 30 sec at 95°C, annealing for 30 sec at varying temperatures as per the primer pairs, and elongation at 72°C for varying lengths of time depending upon the amplicon sizes, followed by 5 min at 72°C. Phusion polymerase was used for all genomic PCRs. For the PCRs in Figure 3B, the 25 μL reaction volume contained 1.0 unit of *Taq* DNA Polymerase (MO267S, New England BioLabs), 1 X ThermoPol Buffer, 0.40 μM of each primer, 0.160 mM of each dNTP, and 0.5 μL of gel extracted DNA fragments as templates. The PCR products were resolved on 1% agarose gels and imaged by ChemiDoc XRS+ system (Bio-Rad Laboratories). Image Lab™ Software 6.0.1 (BioRad Laboratories) was used to quantify band intensities followed by Student’s t test analysis on Microsoft Excel to obtain p-values.

**Table I.**
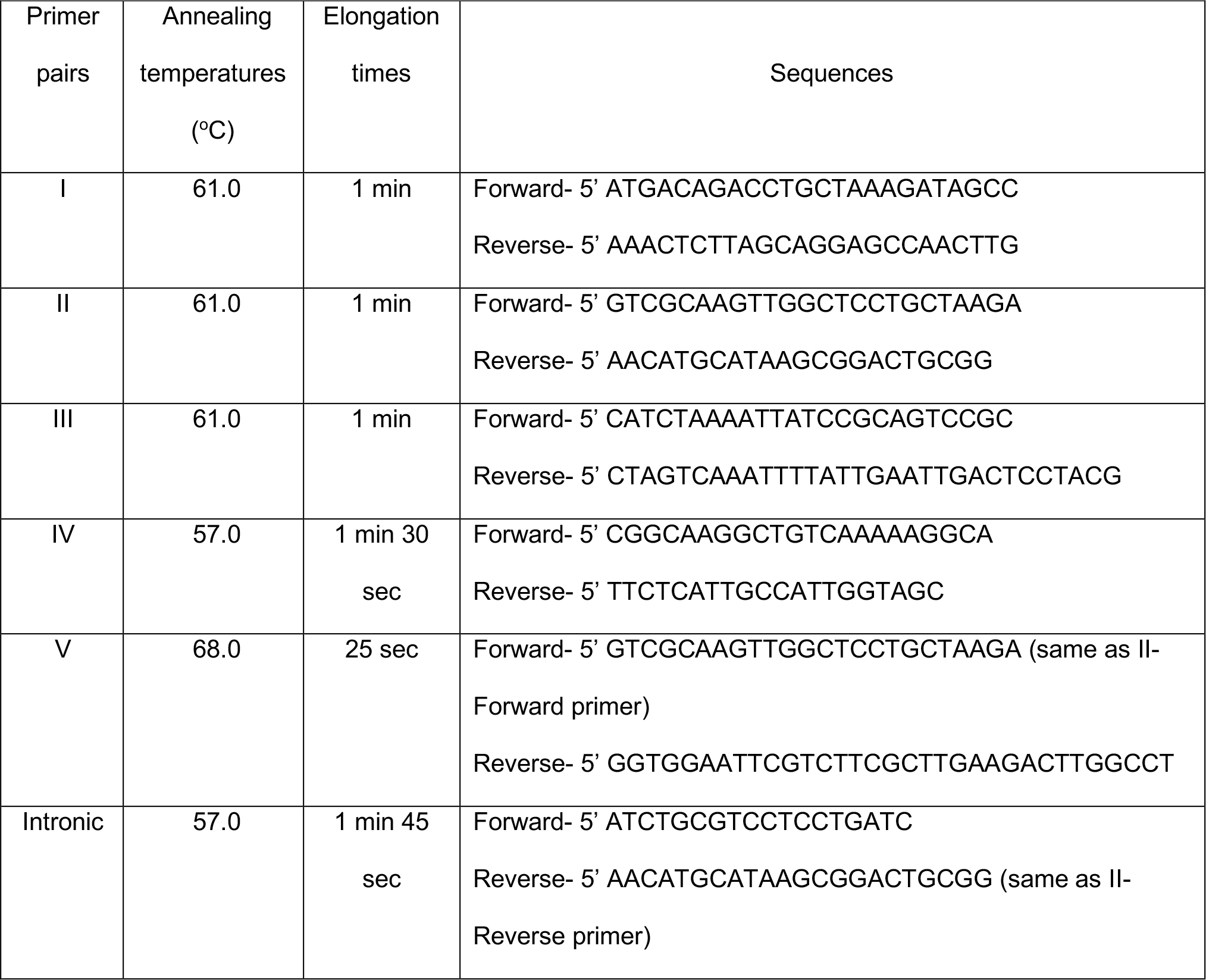
Primer sequences and PCR conditions

### DNA sequencing and sequence analysis

Following phenol-chloroform gel extraction and ethanol precipitation, the PCR products were sequenced with an ABI 3130XL Genetic Analyzer (Applied Biosystems) using BigDye Terminator Cycle Sequencing kit v.3.1. The forward and reverse primers used for PCR amplification (Table I) were used as primers to sequence the PCR products in both forward and reverse directions. Sequences were analyzed and aligned using CLC Sequence Viewer (QIAGEN Bioinformatics).

### Data availability

All fly stocks obtained through Bloomington Drosophila Resource Center can be purchased from the center. *Nopp140* gene deletion line is available upon request. Sequences of all primers used in this study are provided in Table I. Supplemental figures that include raw sequence reads are available in figshare.

## Results

### Repeat sequences within the *Nopp140* second exon

The large central domain of both Nopp140 isoforms in *Drosophila melanogaster* consists of several alternating basic and acidic motifs. This domain is encoded by the second exon of the *Nopp140* gene (Flybase CG7421; FBgn0037137). We found a large segment within this second exon that can be displayed as three distinct but overlapping repeat patterns; we designate the patterns as P”, P’, and P (Figure 1A). The repeat patterns begin respectively at base pairs 874, 900, and 938 with these numbering designations starting at the first base pair of the second exon (Figure 1A). Repeat pattern P’ consists of three tandem repeats of 96 bp each, and repeat pattern P” consists of three 96 bp tandem repeats and a fourth incomplete repeat of 26 bp. On the other hand, repeat pattern P has five tandem repeats of 48 bp flowed by a sixth incomplete repeat of 10 bp.

These repeat patterns contribute to the alternating basic (green) and acidic (yellow) motif pattern within the central domain of both Nopp140 isoforms in *Drosophila melanogaster* (Figure 1B). For instance, the 96 bp repeat sequence in the P’ pattern encodes basic residues (lysine) in its first half and acidic residues (glutamate and aspartate) along with serine residues in the second half: APAKATPAKAIPAKKAASSDDSSSEEEAPKK. These serine residues are likely phosphorylated by casein kinase type II enzymes (Meier 1996) to contribute to the acidic properties of the region.

When comparing the second exon of *Nopp140* from *Drosophila melanogaster* with *Nopp140* genes from other *Drosophila* species (*D. simulans, D. yakuba, D. erecta, D. ananassae, D. pseudoobscura, D. virilis, and D. grimshawi*), we found that only *D. melanogaster* carries the repeating patterns P”, P’ and P. The other species carry roughly one third of the entire repeat segment (Figure 1C). For instance, while *D. melanogaster* and *D. simulans* are closely related species, *D. simulans* lacks 195 bps encoding 65 amino acids, essentially the latter two P’ repeats as compared to *D. melanogaster*. This suggests that a single P’ repeat is functionally sufficient for *Nopp140*-like proteins in the other *Drosophila* species, and that the extra repeats may have duplicated independently in *D. melanogaster*.

### *Nopp140-Long and Nopp140-Short* alleles in *Drosophila melanogaster*

Here we describe a polymorphism in the *Nopp140* gene of *Drosophila melanogaster* that involves the number of P’ pattern repeats described in the previous section. The allele with two repeats is referred to as *Nopp140-Short*, while the allele with three repeats is referred to as *Nopp140-Long*. Both alleles appear to be functional: fly lines homozygous for one or the other allele show no discernible abnormality. We investigated five fly lines (*w^1118^, Oregon-R-C, Canton-S, Daughterless-GAL4*, and *TM3/Et^50^*). For each fly line, we used three PCR primer pairs I, II, and III (Table 1) to amplify the *Nopp140* gene region from genomic DNA extracted from forty adult flies. Together, the three primer pairs covered the entire *Nopp140* gene (Figure 2A). While primer pairs I and III each produced a single PCR product of expected size (1046 bp and 1280 bp, respectively) in all five fly lines, primer pair II generated either one or two PCR products (881 bp and/or 977 bp) depending on the fly line, more specifically, the long or short alleles (Figure 2A). The *w*^*1118*^ fly line produced only the short PCR product of 881 bp, indicating this stock is homozygous for *Nopp140-Short*. *Oregon-R-C, Canton-S, and Daughterless-GAL4* (*Da-GAL4*) fly lines produced the 977 bp PCR product. These lines are therefore homozygous for *Nopp140-Long*. Primer pair II amplified both PCR products of 881 bp and 977 bp from genomic DNA isolated from the *TM3/Et*^*50*^ fly stock. Thus, this stock is heterozygous for the *Nopp140-Long* and *Nopp140-Short* alleles (Figure 2A). Subsequent work with a *Nopp140* gene deletion (He et al., 2015) balanced with *TM3* showed that this *TM3* balancer chromosome contains the *Nopp140-Short* allele (Supplemental Figure S1).

**Figure 2.**
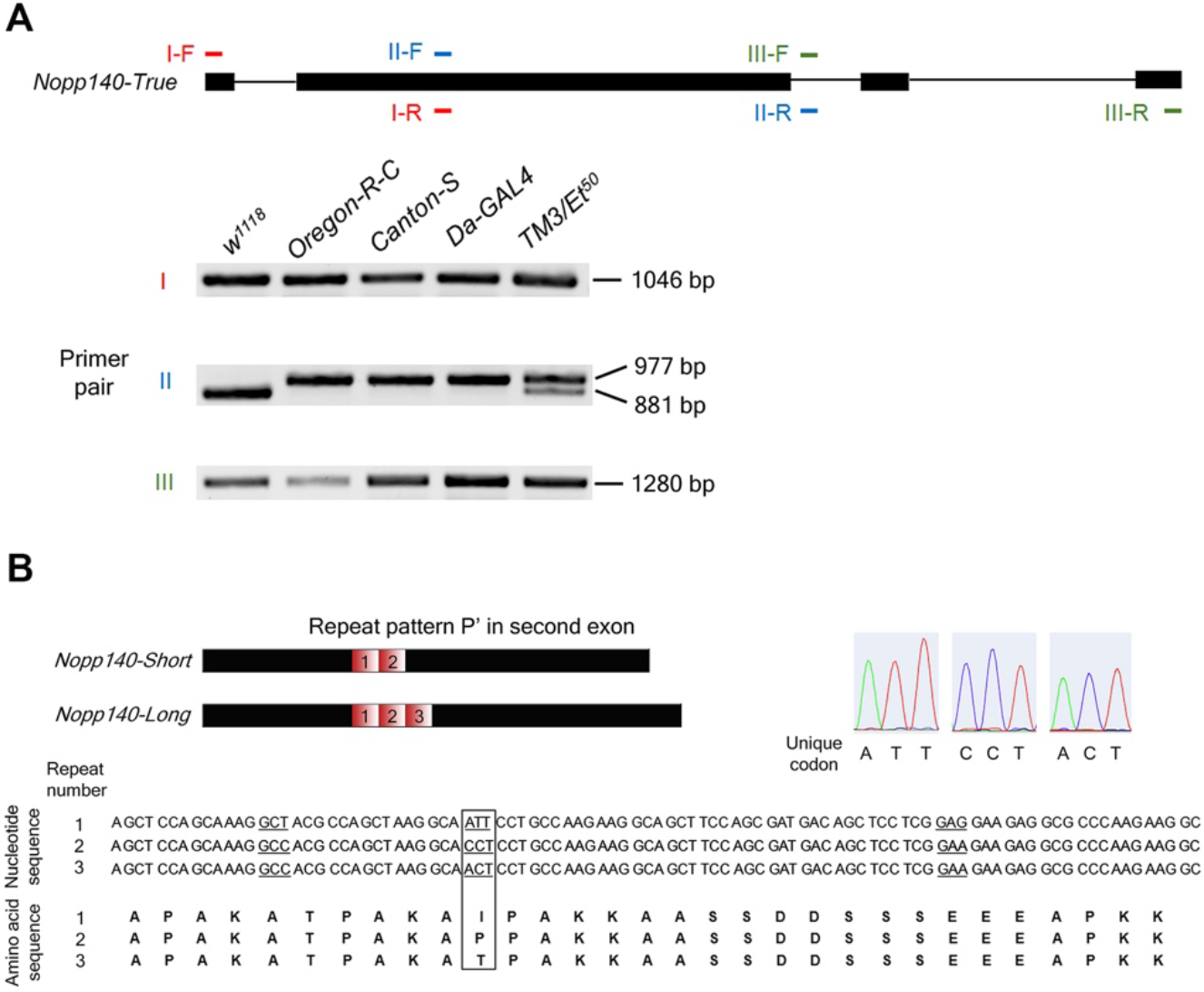
Two *Nopp140* alleles, *Nopp140-Short* and *Nopp140-Long*, exist in *Drosophila melanogaster*. **A** PCR amplification of *Nopp140* genomic DNA region (from start to stop codon for Nopp140-True isoform) in five *Drosophila* fly lines (n=40; 20 adult male flies and 20 adult female flies): *w*^*1118*^, *Oregon-R-C, Canton-S, Daughterless-GAL4 (Da-GAL4)*, and *TM3/Et*^*50*^, using three primer pairs, I, II and III (F = Forward, R = Reverse). **B** *Nopp140-Short* and *Nopp140-Long* alleles have two and three repeats respectively, corresponding to repeat pattern P’, and the repeats are numbered 1, 2 and 3 in the 5’ to 3’ direction on the coding strand. The nucleotide sequences of the three repeats of pattern P’ and the corresponding amino acid sequences for each repeat are shown. The codons at which the three repeats differ are underlined, and within the box are the codons that encode for variable amino acids that are unique to each repeat.

Sequencing the various PCR products obtained from the five fly lines (Figure 2A) revealed that the *Nopp140-Short* allele consists of two P’ repeats (1 and 2), whereas the *Nopp140-Long* allele contains three P’ repeats (1, 2, and 3), all of which are exactly 96 bp in length. Each repeat encodes a 31 amino acid peptide sequence, with functional codons starting with the second nt within the repeat thus leaving 2 nts at the end of the repeat (see Figure 2B).

The nucleotide sequence of the three repeats is identical except at three codons (codons 5, 11, and 25; underlined in Figure 2B). Codon 5 encodes alanine in all three repeats using either GCT or GCC. Codon 25 encodes glutamate in all three repeats using either GAG or GAA. Interestingly, codon 11 encodes amino acids that are unique to each repeat; specifically, repeat 1 uses ATT encoding isoleucine, repeat 2 uses CCT encoding proline, and repeat 3 uses ACT encoding threonine (boxed enclosure in Figure 2B; the sequencing chromatograms for these unique codons are provided). Thus, we can distinguish among the three repeats of the P’ pattern using these particular codons. While sequencing the PCR primer pair II products, we simultaneously sequenced the PCR products obtained with primer pairs I and III; the sequences for these PCR products were identical among all five fly lines examined (Supplemental Figure S2).

### Preferential amplification of *Nopp140-Long* allele in genomic PCRs

In our PCR analyses of fly lines heterozygous for the *Nopp140-Long* and *Nopp140-Short* alleles, such as in *TM3/Et*^*50*^ or *TM3/Sb*, we consistently observed the long PCR product in greater abundance compared to the short product. We used primer pairs II and IV to amplify short and long products from the*TM3/Et*^*50*^ fly line (see Figure 2A and Figure 3A, respectively), and primer pair IV to amplify short and long products from the *TM3/Sb* fly line (Figure 3A). We had similar observations using genomic DNA extracted from the F1 progeny (*Da-GAL4/+*, n=40) using primer pair IV (Figure 3A). These F1 progeny were obtained from a cross between the *Da-GAL4* stock (homozygous for *Nopp140-Long*) and the *w*^*1118*^ stock (homozygous for *Nopp140-Short*). The *Da-GAL4* transgene resides on the third chromosome along with *Nopp140-Long*. Since the *Da-GAL4* transgene is marked with *mini-white^+^*, we can track this particular chromosome in subsequent crosses. We obtained the same unequal product abundance between long and short products when we performed single fly PCR analyses. We used primer pairs IV and V to amplify genomic DNA extracted from single *Da-Gal4/+* F1 adult male flies (Figure 3B). The long PCR product was always in greater abundance compared to the short product.

**Figure 3.**
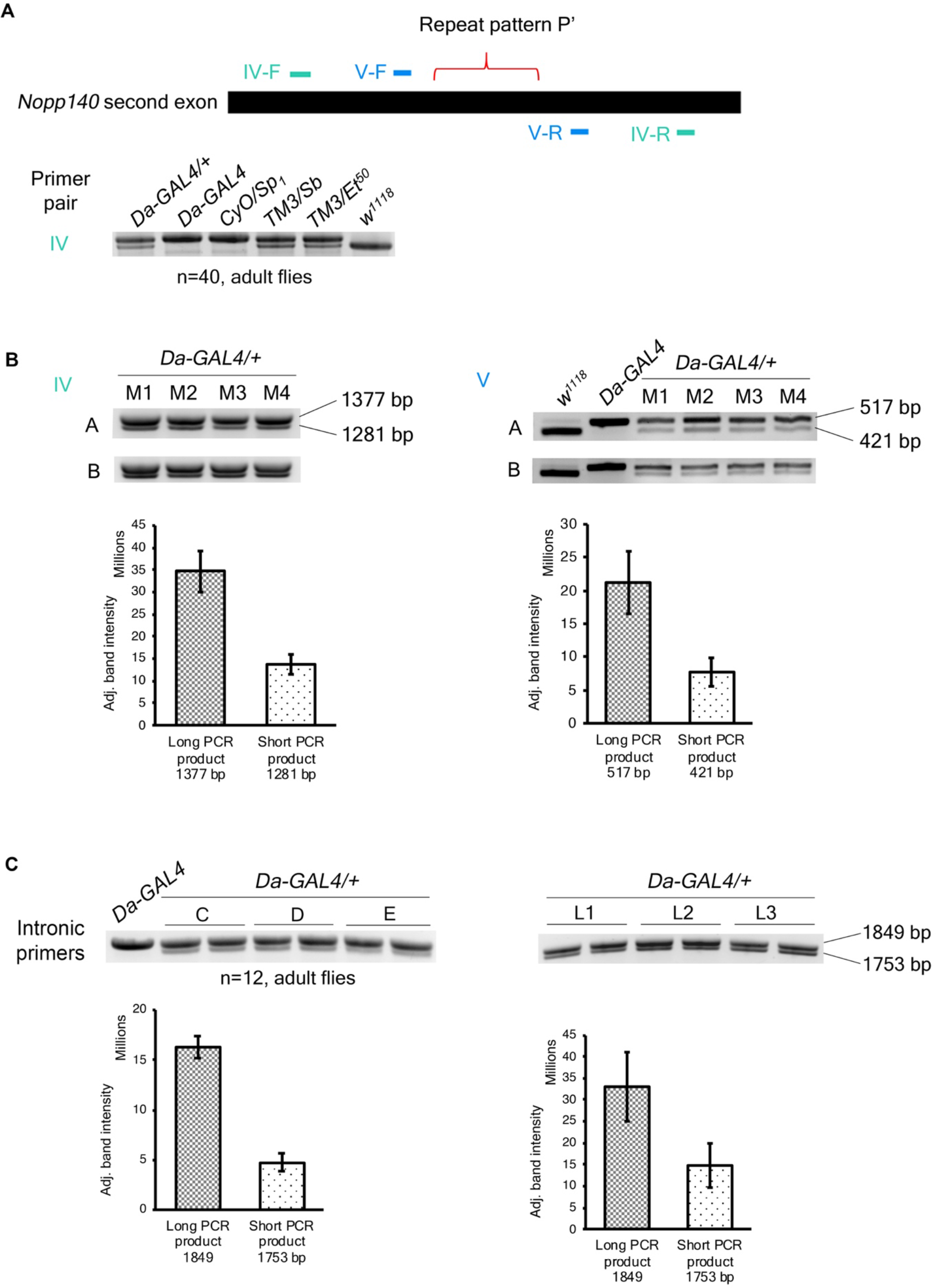
*Nopp140-Long* allele is preferentially amplified in genomic PCRs. **A** PCR amplified *Nopp140* second exon containing repeat pattern P’ in different *Drosophila melanogaster* fly lines (n=40; 20 adult male flies and 20 adult female flies) using two different primer pairs, IV and V (F = Forward, R = Reverse). Primer annealing sites on the *Nopp140* second exon are indicated in the schematic diagram. **B** Primer pairs IV and V were used to amplify the repeat pattern P’ from F1 heterozygotes (*Da-GAL4/+)* obtained from reciprocal crosses (A and B) between *Da-GAL4* and *w*^*1118*^ flies. Two PCR products of 96 bp difference were amplified indicating presence of both long and short *Nopp140* alleles. M1-M4 are PCR samples in which genomic DNA extracted from single adult male flies was used as templates. Quantifications of the band intensities of the long and short PCR products for the single male genomic PCRs are included. An unpaired one-tailed t-test with unequal variance was performed; p-values were 2.1E-07 and 4.6E-06 for PCRs with primer pairs IV and V respectively). **C** PCR amplified *Nopp140* second exon containing repeat pattern P’ in F1 heterozygotes *Da-GAL4/+* (n=12 adult male flies) obtained from three separate crosses (C, D, E) between *Da-GAL4* and *w*^*1118*^ flies, and from individual third instar larvae (L1, L2, L3) obtained from cross B, using an intronic primer pair that spans the entire *Nopp140* second exon. Quantification of the band intensities of the long and short PCR products from the adult male genomic PCR and the single third instar larva genomic PCR are included. An unpaired one-tailed t-test with unequal variance was performed; p-values were 1.5E-09 and 2.4E-4 for PCRs using genomic DNA of twelve adult flies and single third instar larvae respectively.

Since we were dealing with repeat sequences, we considered the possibility that the overabundance of the long product was a PCR artefact. Therefore, we amplified the repeat region with primer pairs that were set further away from the repeats and within the two introns flanking the second exon; increasing distance of the primer annealing sites relative to the repetitive sequence is known to improve amplification of major PCR products and significantly reduce undesired PCR artefacts (Hommelsheim et al. 2014). With these intronic primers, we expected equal amplification of the long and short PCR products from heterozygous flies. However, the long PCR product was still more abundant than the short PCR product when using genomic DNA extracted from both male adult flies (n=12) from three separate crosses, C, D, and E, and from a set of three single third instar larva, L1-3 (Figure 3C).

Based on these results, we considered the possibility of an *in vivo* gene conversion event occurring in the heterozygous flies whereby an insertion of a 96 bp repeat into the *Nopp140-Short* alleles would increase the abundance of the *Nopp140-Long* alleles. This could explain the overabundance of the long PCR product in the heterozygous genomic PCRs. In order to test for this possibility, we amplified the second exon of *Nopp140* from a mixture of genomic DNAs containing equal amounts (50 ng each) from *w*^*1118*^ flies (homozygous for *Nopp140-Short* allele) and *Da-GAL4* flies (homozygous for *Nopp140-Long* allele) using the same intronic primers. This *in vitro* mixing should closely mimic the genomic DNA sample from heterozygous flies, and eliminate the possibility of any gene conversion events. Once again, the long PCR product was more abundant than the short PCR product (Figure 4A) indicating that this is simply a PCR artefact, but one that is consistent and predictable.

**Figure 4.**
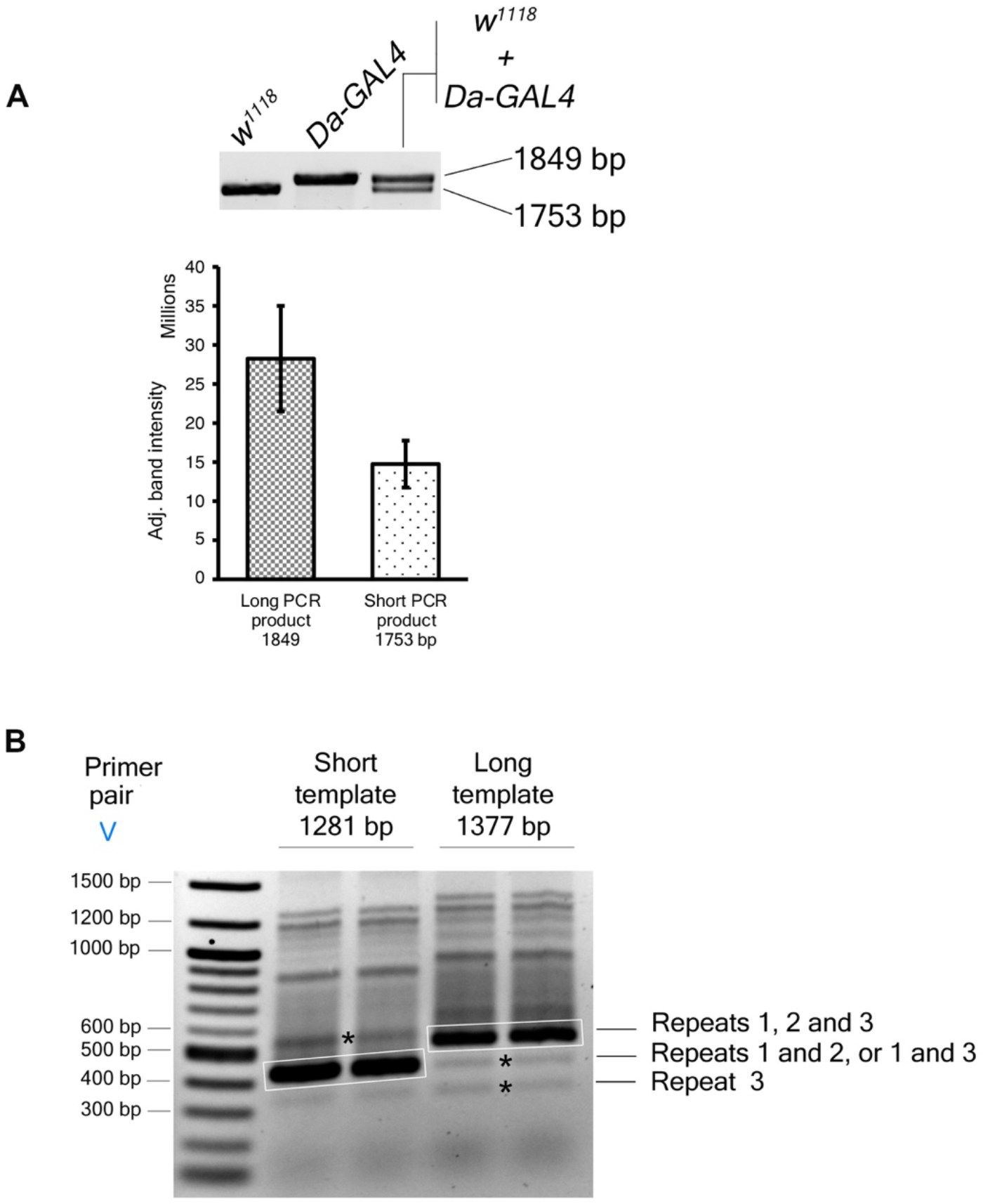
PCR of repetitive DNA sequences generates artefacts. **A** Genomic PCRs using an intronic primer pair and one of the three templates: 50 ng of w1118 genomic DNA (contains only the Nopp140-Short allele), 50 ng of *Da-GAL4* genomic DNA (contains only the Nopp140-Long allele), or a mixture of 50 ng each of *w*^*1118*^ and *Da-GAL4* genomic DNA (contains both Nopp140-Short and Nopp140-Long alleles in equal quantities). Two PCR products of 96 bp difference, 1849 bp and 1753 bp, were amplified among which the longer one was over amplified in PCR with the mixed genomic DNA sample (*Da-GAL4* + *w^1118^*). An unpaired one-tailed t-test with unequal variance was performed for the mixed genomic PCR sample (n=5 reactions); a p-value of 0.0040 was obtained. **B** PCR using primer pair V amplified the repeat pattern P’ from either a 1281 bp short template DNA containing two 96 bp repeats or a 1371 bp long template DNA containing three 96 bp repeats. The bands enclosed in white boxes are the major PCR products of expected sizes, and the undesired fragments that were also sequenced and analyzed in this study are indicated by asterisks.

In summary, we report a PCR phenomenon in which the longer genomic fragment with repeats 1, 2 and 3 was preferentially amplified compared to the shorter genomic fragment with two repeats 1 and 2, despite equal amounts of the *Nopp140-Short* and the *Nopp140-Long* alleles present as templates. We never observed the opposite result where the short product was more abundant than the long product.

### PCR amplification of undesired fragments with repetitive DNA sequences

Studies have shown that amplification of DNA with repetitive sequences can produce an array of undesired fragments with variable number of repeats (Hommelsheim et al. 2014). Indeed, upon amplification of the repeat pattern P’ with primer pair V, PCR products of variable sizes were amplified producing a laddering effect (Figure 4B). There were undesired PCR products both longer and shorter than the major PCR product (boxed in Figure 4B). Surprisingly, when using a short DNA fragment of 1281 bp (the short template) that contained two P’ repeats (1 and 2) as template, sequencing analysis confirmed that one of the higher molecular weight PCR fragments (marked by an asterisk in Figure 4B) contained three P’ repeats: 1, 2, and 3 (in this order 5’ to 3’, each distinguishable by the unique codons described earlier; similar to results shown in Supplemental Figure S4). Likewise, when using the long template (1377 bp) that had three P’ repeats (1, 2, and 3), one of the lower molecular weight PCR products contained two P’ repeats 1 and 2, another had two P’ repeats 1 and 3, and yet another had either repeat 1 or 3 (marked by asterisks in Figure 4B; See Supplemental Figure S3).

## Discussion

Vertebrate *NOCL1* and *TCOF1* genes use multiple exons to encode alternating acidic and basic motifs that repeat to comprise large central domains in their protein products. The *Drosophila Nopp140* gene, however, uses a single large exon to encode a similar large central domain with alternating acidic and basic repeating motifs. We described three repeat peptide sequence patterns within this central domain. The repeats within a single pattern occur in tandem, but the three different patterns overlap with each other (Figure 1A). One of the patterns, P’, consists of 96 bp tandem repeats in *Drosophila melanogaster*, but other closely related *Drosophila* species did not carry the sequence as tandem repeats, suggesting that the repeat region in the *Drosophila melanogaster Nopp140* gene may have originated relatively recently in evolutionary time.

Here we identified length polymorphisms affecting the number of P’ repeats. Various fly lines carry either two or three 96 bp P’ tandem repeats. Hence, we describe the two *Nopp140* alleles as either *Nopp140-Short* (two repeats) or *Nopp140-Long* (three repeats). The individual repeats can be identified by single nucleotide polymorphisms that exist in codon 11 (see Fig 2B); the *Short* allele contains repeats 1 (ATT) and 2 (CCT), and the *Long* allele contains repeats 1 (ATT), 2 (CCT), and 3 (ACT). We described *D. melanogaster* fly lines that are homozygous for either the long allele or the short allele. Balanced fly lines can be heterozygous for both alleles.

### Possible PCR Artefacts

We consistently observed the second exon of *Nopp140-Long* allele was over produced by PCR compared to that of the *Nopp140-Short* allele. The two alleles should be present in equal numbers in the genomic DNA extracted from heterozygous individuals, and hence PCR amplification should theoretically yield equal amounts of PCR products from the two alleles. Initially we considered an *in vivo* gene conversion event in which the *Nopp140-Short* allele is preferentially converted to the *Nopp140-Long* allele as a possible explanation because we found that *w*^*1118*^ genomic PCRs amplified a minor PCR product of the size expected for a long PCR product on certain occasions (Supplemental Figure S4). While we did not carry unbalanced heterozygotes (*Da-GAL4*/*w^1118^*) past two generations to test the stability of the long versus short alleles, we did test different larval tissues to see if amplification of the long allele was restricted to polyploid (midgut) versus diploid (brain) tissues, but we saw no difference; the long allele was always preferentially amplified regardless of the source of genomic DNA.

In a simple mixing experiment, we combined genomic DNA homozygous for *Nopp140-Long* with equal amounts of genomic DNA homozygous for *Nopp140-Short*, and observed the same preferential amplification of the long allele. This *in vitro* mixing experiment refutes an *in vivo* gene conversion event. Confusing the issue however, *w*^*1118*^ genomic DNA is homozygous for *Nopp140-Short* allele containing only tandem repeats 1 and 2, but we identified minor amplicons from *w*^*1118*^ genomic DNA with repeat pattern P’ containing tandem repeats 1, 2 and 3 (Supplemental Figure S4). Likewise, we consistently amplified (and sequenced) a 521 bp PCR product with repeat pattern P’ that contained tandem repeats 1, 2 and 3 (similar to the results shown in Supplemental Figure S4), although the 1281 bp template with repeat pattern P’ contained only tandem repeats 1 and 2.

We have yet to explain this preferential PCR amplification of the long product versus the short product, but we may have an explanation for the amplification of short PCR products with repeats 1 and 2, 1 and 3, or a single repeat 1 (Figure 3B) from the 1371 bp template DNA that contained all three tandem repeats. Studies by others have shown that PCR amplification of repetitive DNA regions can generate undesired fragments of variable sizes containing haphazard combinations of repeats, similar to our observation shown in Figure 3B. PCR products with variable repeats could potentially arise from an initial production of incomplete, single-stranded fragments that then acted as mega-primers in subsequent PCR cycles. These mega-primers would then misalign with the existing templates at the repetitive sites, thereby resulting in an array of PCR fragments of sizes other than the major product (Hommelsheim et al. 2014).

### Implications for Long versus Short *Nopp140* Alleles

Earlier, we showed that partial depletion of Nopp140 in *Drosophila* by siRNA expression resulted in *Minute*-like phenotypes such as deformed wings, legs, and tergites (Cui and DiMario, 2007). These phenotypes were similar to those observed with the various haplo-insufficiency *Minute* mutations in genes encoding ribosomal proteins, and reminiscent of the phenotypes associated with the TCS (Sæbøe-Larssen *et al.*, 1997; Cui and DiMario, 2007). Flies homozygous for either the short allele or the long allele, or flies heterozygous for the two alleles are viable and do not exhibit any of the discernible phenotypes associated with the partial depletion of the Nopp140 isoforms.

While the P’ repeat polymorphism does not seem to have a significant impact on the core functions of Nopp140, the protein’s central domain is generally considered to be an unstructured region. Recent studies have revealed the important role of inherently disordered proteins (IDPs) in promoting phase separation to form subcellular membrane-less compartments or assemblages (Toretsky and Wright, 2014). Nopp140 resides in the nucleolus and the Cajal bodies; both are membrane-less compartments within nuclei. The large central domain of Nopp140 likely contributes to a myriad of protein-protein interactions and protein-RNA interactions that occur in these sub-nuclear compartments. Future work should establish how this unstructured central domain of Nopp140 may be involved in these interactions and perhaps the phase-separations of the nucleolus and Cajal bodies.

